# Genome-wide association and environmental suppression of the mortal germline phenotype of wild *C. elegans*

**DOI:** 10.1101/2023.05.17.540956

**Authors:** Lise Frézal, Marie Saglio, Gaotian Zhang, Luke Noble, Aurélien Richaud, Marie-Anne Félix

**Author notes:** : Co-first authors (Lise Frézal, Marie Saglio).

## Abstract

The animal germline lineage needs to be maintained along generations. However, some *Caenorhabditis elegans* wild isolates display a mortal germline phenotype, whereby the lineage becomes sterile after several generations at 25°C. We used a genome-wide association approach to study the genetic basis for this phenotype in *C. elegans* populations. We detected a significant peak on chromosome III around 5 Mb, which was confirmed using introgression lines. These results indicate that a seemingly deleterious genotype is maintained at intermediate frequency in the species. Environmental rescue is a likely explanation and we indeed find that naturally associated bacteria and microsporidia suppressed the phenotype. The tested bacteria also suppressed the temperature-sensitive mortal germline phenotype of mutants in small RNA inheritance (*nrde-2*) and histone modifications (*set-2*). Even *Escherichia coli* strains of the K-12 lineage suppressed the phenotype compared to B strains. By shifting a strain cultured on *E. coli* K-12 back to *E. coli* B, we found that *C. elegans* can keep over several generations the memory of the suppressing conditions. Thus, the mortal germline phenotype of wild *C. elegans* is lin part revealed by laboratory conditions and may represent variation in epigenetic inheritance and environmental interactions. This study also points to the importance of non-genetic memory in the face of environmental variation.

## Introduction

The animal germline is an immortal cell lineage that persists through organismal generations. The nematode *Caenorhabditis elegans* displays a short generation time of about three days under laboratory conditions, allowing studies of the conditions for germline immortality. Indeed, Ahmed and Hodgkin screened for mutants of the N2 reference strain with a "mortal germline" (Mrt) phenotype, defined as a progressive onset of sterility across generations (Ahmed and Hodgkin 2000; Smelick and Ahmed 2005).

Among the genetic loci thus defined, a subset displayed a temperature-sensitive (ts) Mrt phenotype, even as null alleles (Sakaguchi et al. 2014). Over the last 10 years, a number of loci were shown to display this ts-Mrt phenotype. The corresponding mutations so far affect nuclear small RNA pathways, small RNA amplification and histone modifications in the germline, relating the ts-Mrt phenotype to transgenerational epigenetic inheritance (Buckley et al. 2012; Robert et al. 2014; Spracklin et al. 2017; Weiser et al. 2017; Manage et al. 2020; Wan et al. 2021).

Epigenetic inheritance of gene expression states operates in *C. elegans* through small RNAs and histone modifications (Ketting and Cochella 2021; Frolows and Ashe 2021; Grishok 2021; Seroussi et al. 2022; Phillips and Updike 2022; Quarato et al. 2022). Briefly, small RNAs bound to specific Argonaute proteins enter the nucleus and induce histone modifications such as repressive H3K9me3 or H3K27 methylation at the corresponding locus, thereby silencing gene expression (Gu et al. 2012; Mao et al. 2015). Small RNAs corresponding to germline-expressed genes are amplified in perinuclear germ granules (Phillips et al. 2012; Wan et al. 2018; Ishidate et al. 2018; Uebel et al. 2018; Wan et al. 2021), allowing for persistence of silencing for several generations. A subset of mutants in nuclear RNAi (such as *nrde-2*) or in germline nuclear Argonaute genes (such as *hrde-1*) display the ts-Mrt phenotype (Burton et al. 2011; Buckley et al. 2012). In addition, mutants in histone methyltransferases, such as *set-2/SET1* (encoding the histone H3K4me3 methyltransferase), display the ts-Mrt phenotype. A balance of histone modifications appears necessary to maintain germline pluripotency and DNA integrity (Katz et al. 2009; Alvares et al. 2014; Robert et al. 2014; Herbette et al. 2017; Robert et al. 2020; Saltzman et al. 2018; McMurchy et al. 2017). Thus, the ts-Mrt phenotype may be an indication of variation in gene silencing mechanisms.

We previously reported that, surprisingly, multiple *C. elegans* wild isolates displayed the ts-Mrt phenotype (Frézal et al. 2018). This phenomenon is interesting for two reasons: first, a sterility phenotype is an odd one to find in a wild population; second, this finding points to possible natural genetic variation in transgenerational epigenetic inheritance. In a first study, we focused on a genetic cross between two isolates of contrasting Mrt phenotype (MY10 and JU1395), which identified a causal deletion in the *set-24* gene underlying the strong Mrt phenotype of MY10. This gene encodes a SET-domain protein, likely involved in producing or reading posttranslational histone modifications. The *set-24* deletion allele is rare across the species (Frézal et al. 2018). Therefore, its evolutionary importance is difficult to assess.

Saber et al. recently reported on the frequency of the Mrt phenotype in wild *C. elegans* isolates and its low mutational variance using mutation accumulation lines. They suggested that the phenotype - or the underlying genetic polymorphisms - may be under balancing selection (Saber et al. 2022). We here asked 1) whether we could detect a common polymorphism underlying the Mrt phenotype of *C. elegans* wild isolates; 2) whether the phenotype could be suppressed by natural environmental conditions.

We first perform a genome-wide association of the Mrt phenotype on over 100 *C. elegans* wild isolates and detect a signal on chromosome III at a genomic location close to 5 Mb. This locus could be confirmed using introgression lines. To explain this common occurrence of a deleterious phenotype, we isolated natural populations of *C. elegans* with their associated microbes. The strains deprived of their natural microbiota exhibited a Mrt phenotype, but showed a strong rescue when reassociated with isolated bacteria or microsporidia. *E. coli* strains of the K-12 lineage also suppressed the Mrt phenotype compared to B lineage strains. We report that the Mrt phenotype of *nrde-2*, *set-2* and *set-24* mutants is also suppressed by bacterial associations, and is thus revealed by standard, simple laboratory conditions. Furthermore, we find that a wild *C. elegans* strain can keep during multiple generations the memory of the suppressing bacterial strain on which their ancestors were cultured at 25°C.

## Results

### Genome-wide association of the mortal germline phenotype

We measured the mortal germline (Mrt) phenotype of 132 worldwide *C. elegans* isolates at 25°C, using the assay in Frézal et al. (2018). The phenotype was coded as a quantitative trait, which we call the Mrt value, defined as the mean number of generations to sterility.

The 25°C Mrt phenotype was found to be common among *C. elegans* wild isolates (Figure 1B). The population structure of *C. elegans* includes highly divergent strains from the Pacific region and a less structured population on the Eurasian, African and American continents (Crombie et al. 2019; Lee et al. 2021) and strains in both groups were found to exhibit a Mrt phenotype (Tables S1 and S2).

**Figure 1.**
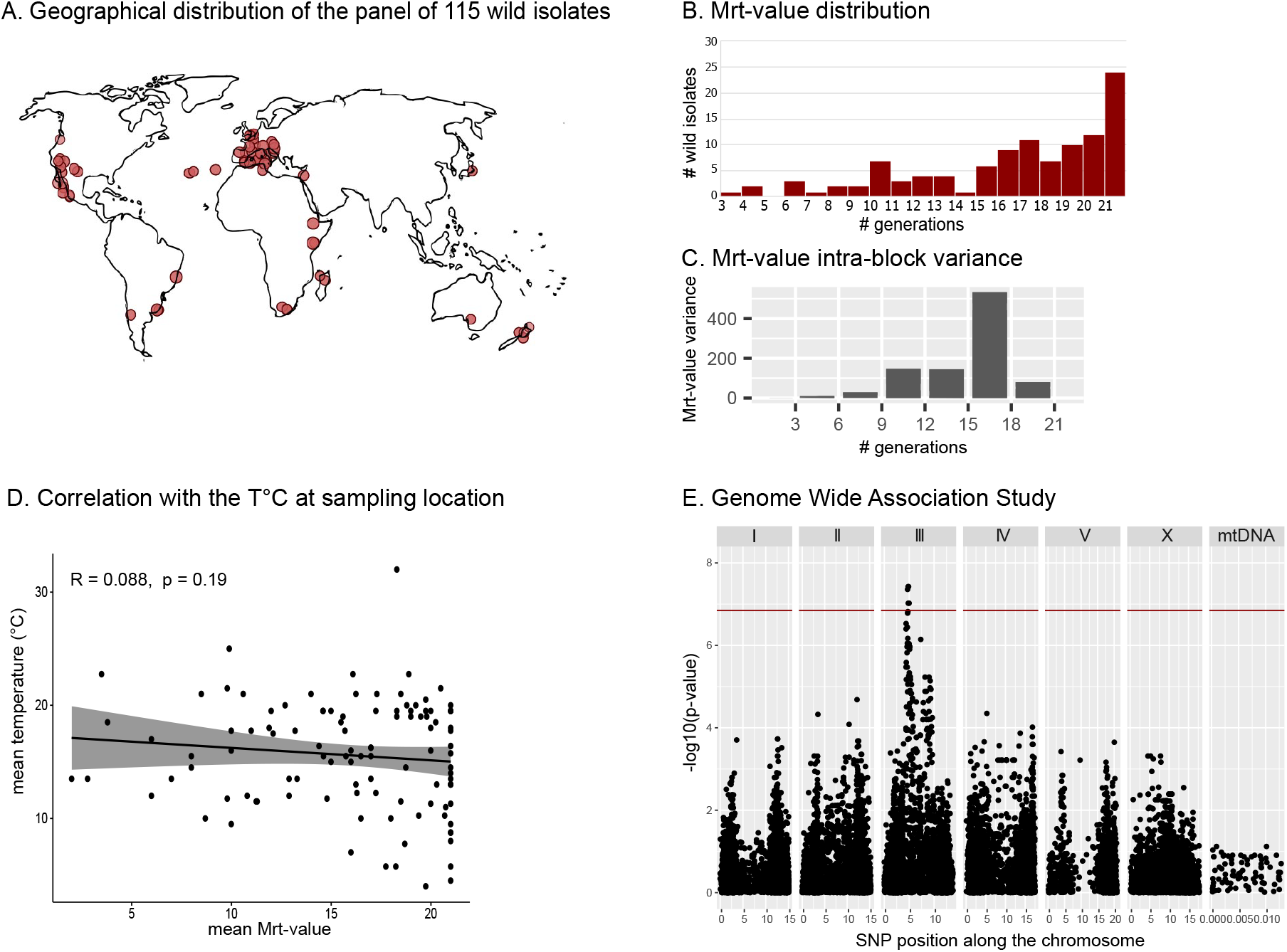
Association of the mortal germline (Mrt) phenotype with a locus on chromosome III in a panel of 115 *C. elegans* isolates. (A) Geographical distribution of the panel of 115 wild isolates, excluding the divergent Pacific strains. (B) Phenotypic distribution of the mortal germline phenotype in the panel, assayed as in (Frézal et al. 2018). The phenotype is expressed in number of generations to sterility at 25°C, until generation 20. Fertility at this last generation is plotted as generation 21. (C) Intra-block variance, plotted as a function of the Mrt-value mean. The Mrt-values are binned in intervals of three generations. (D) Absence of correlation between the Mrt-value and temperature at the sampling location. (E) Genome-wide association. The plot represents the significance value of the association between the tested single-nucleotide polymorphisms (SNPs) plotted along the six chromosomes and mitochondrial DNA (MtDNA). The red line indicates the significance threshold at p<0.05. See Table S3 for fine mapping with the full set of SNPs.

In the restrictive set of 115 strains shown in Figure 1, the mean Mrt-values of strains ranged from sterile at 3 generations to still fertile after 20 generations at 25°C, with a skewed distribution toward high values (Figure 1B). The variance among replicates was low for strong Mrt values and high at intermediate trait values (Figure 1C; Table S2). Given this distribution, we analyzed the data using generalized linear mixed models (cf. Material and Methods). The Mrt values were obtained from different experimental blocks, including different experimenters and laboratories. A significant block effect was detected, which was taken into account in the analysis. For GWAS mapping, we included data collected in Saber et al. (2022) and used either the full strain set, or restricted sets without the divergent Pacific area strains.

Using GWAS with the restricted set of 115 strains, we found that variation in the number of generations to sterility was significantly associated with genetic variation in a region of chromosome III located between 4 and 6 Mb (Figure 1E). The larger strain set including divergent Pacific region strains also gave a significant peak on chromosome III between 4 to 8 Mb (Figure S1). The association was stronger and the peak narrower with the restricted set of isolates, in line with the fact that the additional strains are genetically divergent and practically isolated from the continental recombinant pool (Crombie et al. 2019; Lee et al. 2021).

### Introgression Lines confirm the association with chromosome III

To test the causal effect of chromosome III, we produced chromosome introgression lines between parents of contrasted phenotypes. We chose JU775 as parent with a Mrt phenotype because this isolate carries the intermediate-frequency chromosome III alleles identified in the GWAS analysis. We introgressed the chromosome III of JU775 into the N2 genetic background using repeated backcrosses associated with genotyping, and tested the Mrt phenotype of parents and introgression lines.

The strains with the N2 genetic background could be propagated in the laboratory for over 20 generations at 25°C (Figure 2A - this included SX461 that carries a here irrelevant single-copy *gfp* transgene). By contrast, the strains with the JU775 genetic background were sterile before 8 generations at 25°C (Figure 2A). The strains carrying the JU775 chromosome III in the N2 genetic background, JU3194 and JU3195, showed a stronger Mrt phenotype compared to the non-Mrt parent (Figure 2A). The Mrt values of the chromosome III introgression lines were intermediate between the parental values, indicating that polymorphisms on other chromosomes must explain an additional part of the difference between JU775 and N2.

**Figure 2.**
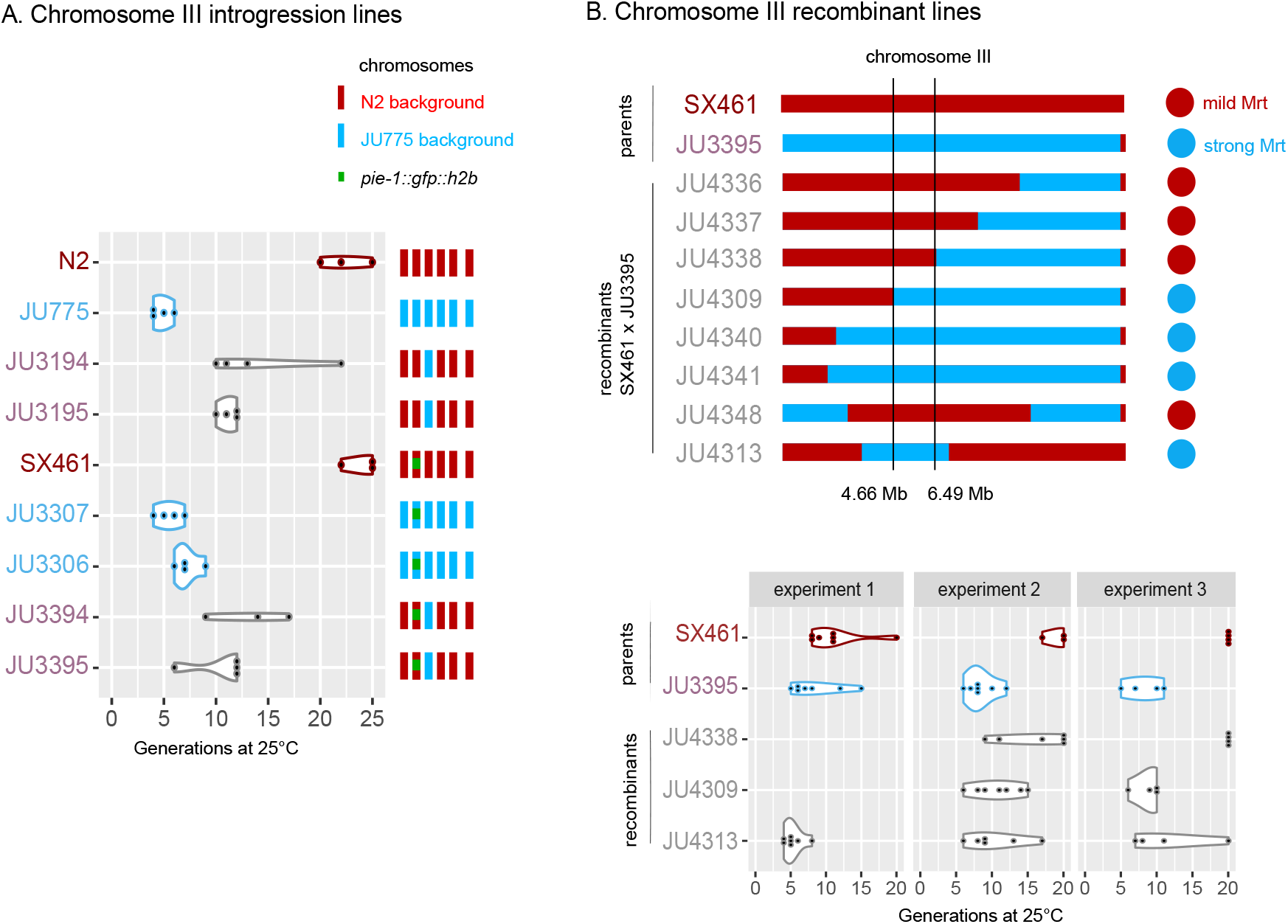
Laboratory crosses confirm the presence of a quantitative trait locus on chromosome III at ca. 5 Mb. (A) Mrt phenotypes of the chromosome III substitution lines. In red is indicated the N2 genetic background (non-Mrt), in light blue the JU775 genetic background (Mrt), and in violet the introgression lines. Some strains carry an irrelevant *gfp* transgene, indicated as a green Both chromosome III and the remaining background had a significant effect by ANOVA on Poisson GLMs (each at p<10^-7^). About 2/3 of the difference in Mrt phenotype is accounted by chromosome III. (B) Mrt phenotypes of the chromosome III recombinant lines. Top panel: schematic of genotypes of the recombinants between JU3395 and SX461, with in blue JU775 alleles and in red N2 alleles. Phenotyping in multiple experiments is shown in Table S2 and schematized on the right in a binary fashion with a blue circle showing a strong Mrt phenotype, a red circle otherwise. Bottom panel: Mrt phenotyping of the parents and a subset of recombinant lines, in three independent experiments. The JU3395 parent and the JU4309 and JU4313 recombinants exhibit a strong Mrt phenotype, while the SX461 parent and the JU4338 recombinant have a mild and more variable Mrt phenotype.

We then screened for recombinants after further crossing the chromosome III introgressed line JU3395 to the N2 genetic background. The recombinant lines were assayed for the Mrt phenotype in several experimental blocks (Table S2). A first group of recombinants displayed a strong Mrt phenotype, comparable to that of the parent JU3395. This group is composed of the recombinants JU4309, JU4340, JU4341 and JU4313. These strains share JU775 alleles in a 4.66-6.49 Mb region on chromosome III (Figure 2B). Therefore, this region is sufficient to give a strong Mrt phenotype. This removed as candidate loci some highly associated alleles on the left side of the GWAS interval (in grey in Table S3). A second group of recombinants that did not include this interval showed a mild Mrt phenotype and thus exhibited more variance, both across replicates of a given experiment, and across experiments (indicated as mild Mrt on Figure 2B; see Table S2). Note that the SX261 (N2 background) parent also showed variation among experimental blocks. Our focus was on the GWAS peak, thus we did not investigate further whether a second locus present in JU775 on the right of chromosome III may have a smaller effect.

Altogether, these results show that a polymorphism located between 4.66 and 6.49 Mb on chromosome III strongly affects the Mrt phenotype in a cross between N2 and JU775. This genomic position is in agreement with the GWAS peak and implies the existence of an intermediate-frequency allele at the species level that causes multigenerational sterility at 25°C in laboratory conditions.

### Naturally associated gut bacteria or microsporidia can rescue the Mrt phenotype of *C. elegans* wild isolates

The Mrt phenotype is intrinsically deleterious since it leads to sterility of the lineage, raising the question of how such polymorphisms can be maintained in natural populations. Temperature fluctuations may in part explain the suppression (Frézal et al. 2018). However, the phenotype was not found to be correlated with the temperature at isolation (Figure 1D) and some isolates with a particularly strong Mrt phenotype in the laboratory, such as JU775, were isolated under warm conditions. We therefore asked whether other ecologically relevant environmental features could affect the Mrt phenotype.

*C. elegans* is generally cultured in the laboratory using *E. coli* OP50 as a food source, yet it encounters a variety of other microbes in the wild (Samuel et al. 2016; Dirksen et al. 2016; Schulenburg and Félix 2017). In the course of *C. elegans* collections from natural samples, we had noticed that the Mrt phenotype commonly appeared after bleaching the culture, a treatment that kills the associated microbes. We thus tested the impact of naturally associated microbes on the phenotype.

To this end, we freshly isolated new *C. elegans* strains from wild sources together with their microbial associates. We assayed the Mrt phenotype of the strains with their associated microbial fauna, or on *E. coli* OP50 alone after a bleach treatment (Figure 3A). The freshly isolated strains JU3224 from New Zealand and JU4134 from Santeuil could be propagated for more than 20 generations at 25°C with their associated microbes. However, after bleaching a culture on *E. coli* OP50, these two strains exhibited a Mrt phenotype at 25°C (Figure 3B,C).

**Figure 3.**
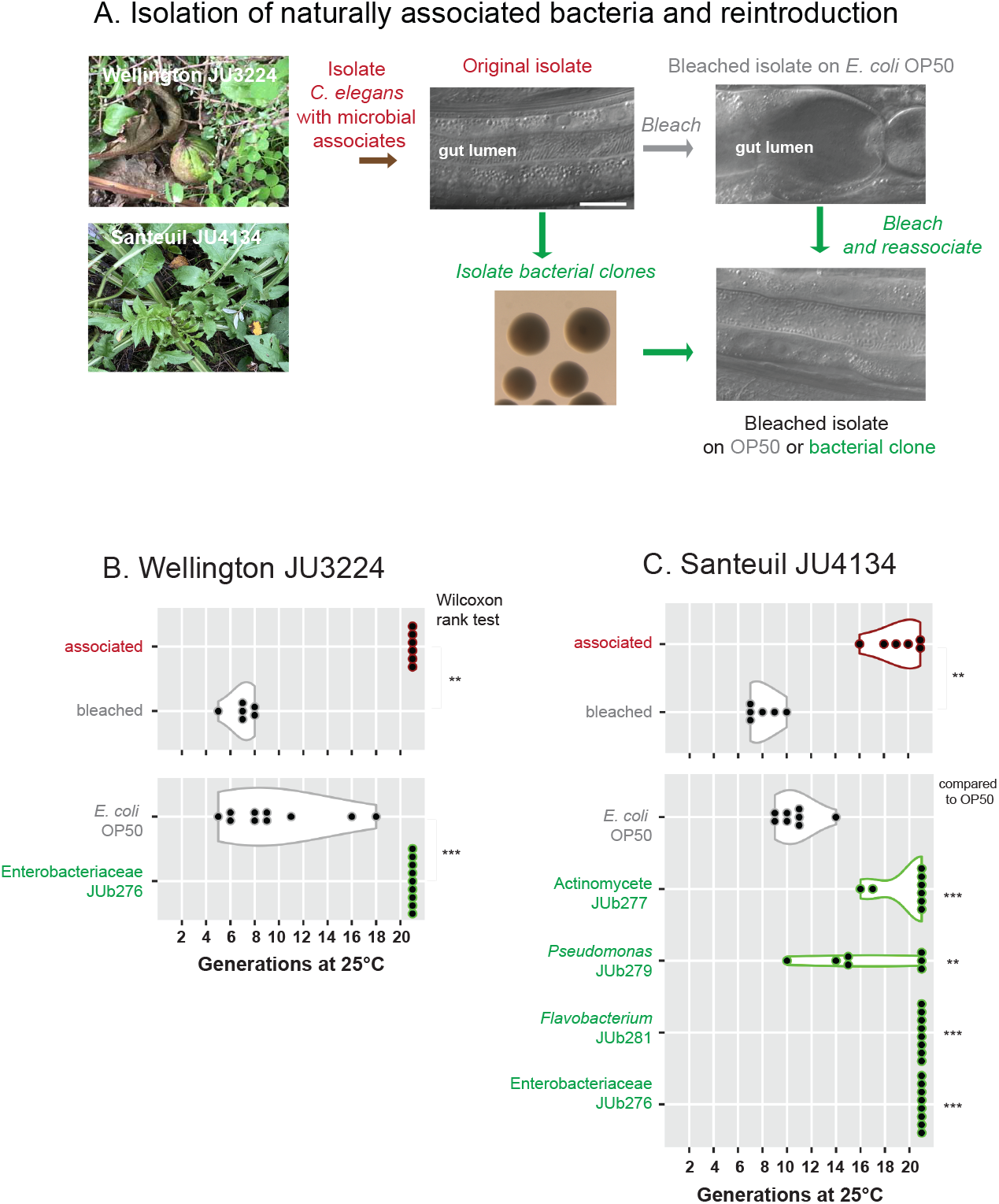
Suppression of the Mrt phenotype of newly isolated *C. elegans* by their associated bacteria. (A) Schematic representation of the experiment, with color coding as in the subsequent panels. *C. elegans* was collected from rotting fruit (JU3224) or stem (JU4134) and bacterial clones isolated from the culture. Nomarski pictures show the presence of associated bacteria in the intestinal lumen of the nematode in the original culture and on the isolated bacteria (here JU3224 on JUb276), but none or few when cultured with *E. coli* OP50. Bar 10 (m. (B) Mrt phenotype of the *C. elegans* isolate JU3224 in the presence of its original microbial associates (brown), on *E. coli* OP50 (grey) or reassociated with its own bacteria JUb276 (green). (C) Mrt phenotype of Santeuil JU4134 in the presence of its original microbial associates (brown), on *E. coli* OP50 (grey) or reassociated with isolated bacteria from its original culture or with JUb276 (green). The experiments of the bottom graphs of panels B and C were performed in parallel. Wilcoxon rank tests: *** for p<0.001; ** p<0.01; n.s.: p>0.05.

From the JU3224 culture, we isolated a *Lelliottia* strain (JUb276) and from the JU4134 culture several bacteria, including an Actinomycete (JUb277), a *Pseudomonas* (JUb279), and a *Flavobacterium* (JUb281). We performed a new assay after reassociating the *C. elegans* strain with either *E. coli* OP50 or the isolated microbes, in groups or singly. The reassociation of single isolated bacterial clones to the bleached wild isolate partially or fully rescued the Mrt phenotype (Figure 3B,C, Figure S2). *Lelliottia* JUb276 from the New Zealand strain could rescue the Mrt phenotype of the bleached Santeuil strain (Figure 3C). This observation suggests a common mechanism of interaction between these bacteria and *C. elegans* that rescued the latter’s sterility.

Microsporidia are obligate intracellular parasites related to fungi. The microsporidia *Nematocida parisii* and *N. ausubeli* are commonly found associated with *C. elegans*, colonizing its intestinal cells (Figure 4A) (Troemel et al. 2008; Zhang et al. 2016). We assayed the effect of their infection on the Mrt phenotype of JU775 and JU2526; the latter isolated in association with *N. ausubeli* (Zhang et al. 2016). These two strains exhibited a strong Mrt phenotype when cultured on *E. coli* OP50 without microsporidia. For both strains, the infection by either species of microsporidia strongly delayed the onset of sterility at 23°C (Figure 4B). Thus, rather than solely acting as a detrimental parasite, microsporidia conferred a context-dependent beneficial effect on their host. In addition, it is worth noting that they interfered with a multigenerational phenotype while infecting the soma.

**Figure 4.**
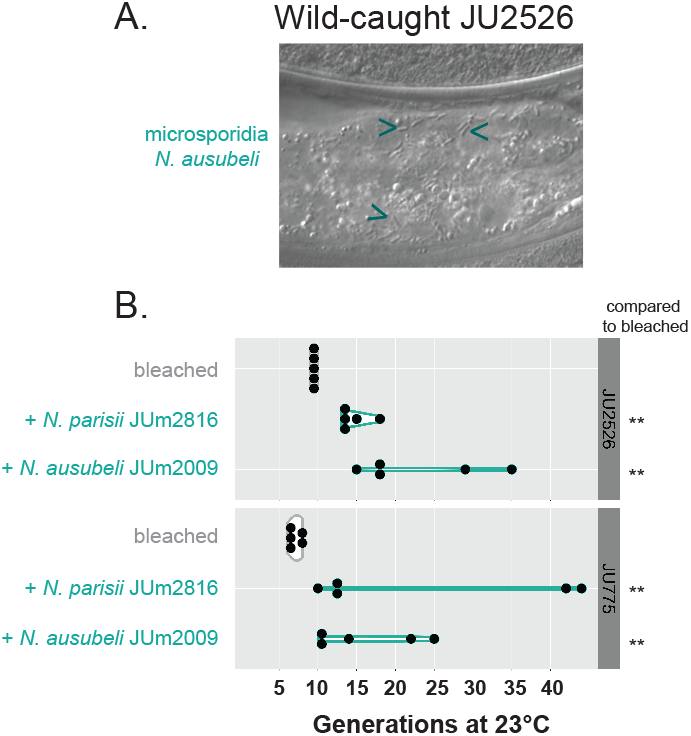
Infection by microsporidia suppresses the Mrt phenotype of *C. elegans* wild isolates. (A) JU2526 was originally associated with microsporidia *Nematocida ausubeli* (Zhang et al. 2016). Arrows indicate microsporidian spores in intestinal cells. Bar 10 μm. (B) The Mrt phenotype of the *C. elegans* isolates JU2526 and JU775 is suppressed by infection with two *Nematocida* strains at 23°C. Wilcoxon rank tests: ** p<0.01.

As we previously reported, starvation or passage through the dauer diapause are common in the wild for *C. elegans* (Barrière and Félix 2005; Félix and Duveau 2012). Such treatments were reported to affect multigenerational phenotypes (Simon et al. 2014; Jobson et al. 2015; Demoinet et al. 2017). We tested whether they could affect the Mrt phenotype of the wild isolate JU775, and did not detect a significant effect in our conditions (Figure S3).

Altogether, the results using naturally associated bacteria and microsporidia highlight the central role of microbial interaction on the phenotype of the host and the effect of infection of somatic tissues on the germline, thus breaching the Weismann barrier.

### *E. coli* K-12 derivatives rescue the Mrt phenotype of *C. elegans* wild isolates compared to B lineage strains

Metabolic differences among *E. coli* strains of relevance to *C. elegans* nutrition are well documented (Yen and Curran 2016; Zhang et al. 2017; Neve et al. 2020). We tested whether a different laboratory strain of *E. coli* could rescue the Mrt phenotype compared to the standard *E. coli* OP50 strain, a B-type strain (Brooks et al. 2009). Indeed, the K-12 strain MG1655 rescued the Mrt phenotype. The rescue was quantitative and depended on the strength of the Mrt phenotype. For the *C. elegans* JU4134 strain with an intermediate Mrt value (around 8-10 generations on OP50 at 25°C), the rescue by *E. coli* MG1655 or the naturally associated bacteria JUb276 and JUb277 was nearly complete (>20 generations for most replicates) (Figure 5A). For the JU775 strain with a strong Mrt phenotype (mean of 4 generations before sterility), the rescue at 25°C was significant but partial (Figure 5A). This partial rescue was likely the consequence of its stronger phenotype and not of a different interaction with the bacterium: the rescue of JU775 was indeed complete at 22°C, a temperature at which the severity of the Mrt phenotype on OP50 was similar to that of the JU4134 strain at 25°C (Figure 5B). Similarly, the Mrt phenotype of the MY10 strain used in (Frézal et al. 2018) was rescued at 22°C (Figure 5C).

**Figure 5.**
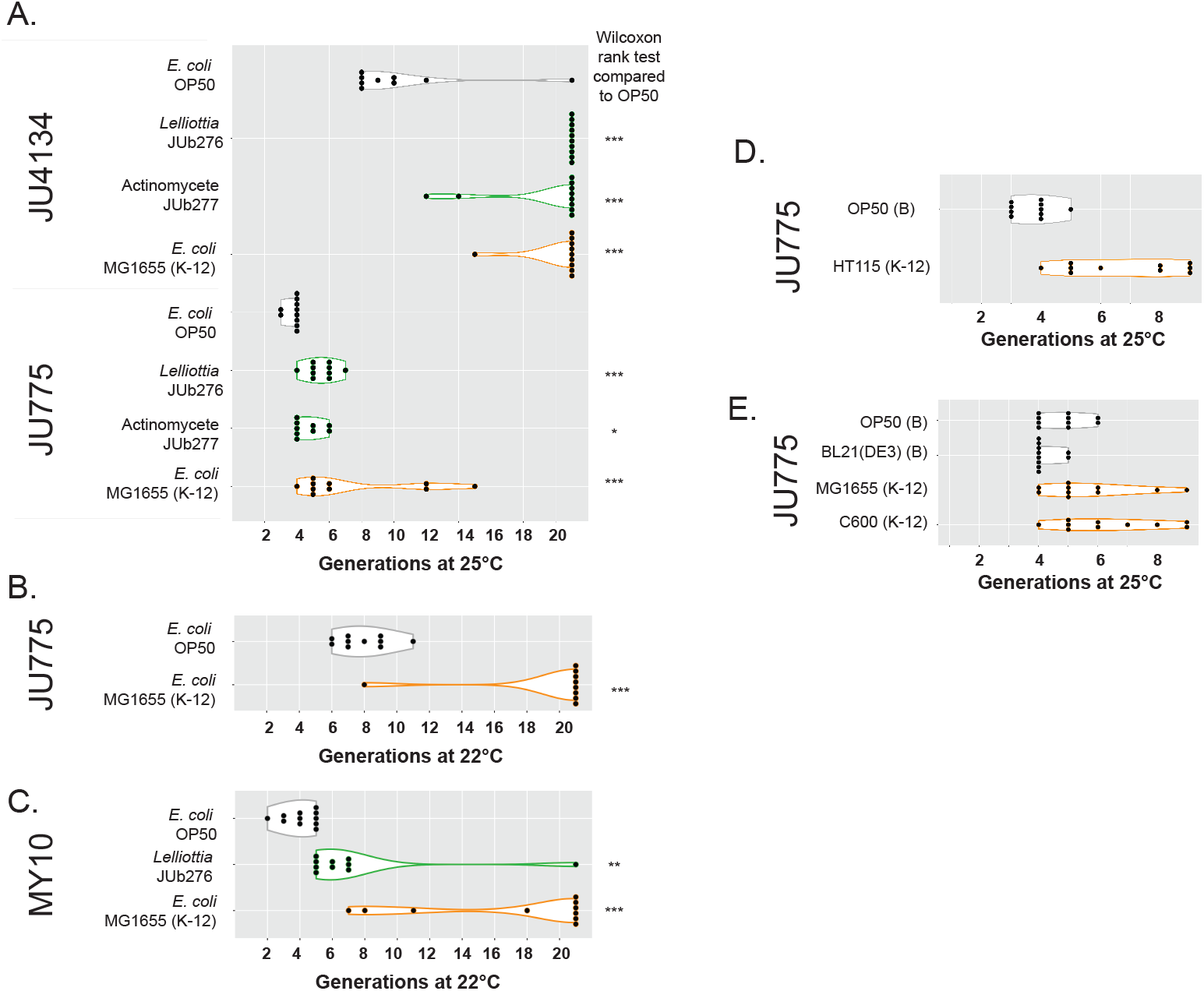
Suppression of mild and strong Mrt phenotypes by naturally associated bacteria and by E. coli K-12 strains. (A-C) The Mortal germline phenotypes of JU4134, JU775 and MY10 are at least partially suppressed by JUb276, JUb277 (natural associates, green) and *E. coli* MG1655 strain (orange). The phenotyping assay was performed at 22°C in (B,C). (D,E) *E. coli* K-12 lineage (orange) but not B lineage strains (grey) partially suppress the Mrt phenotype of *C. elegans* wild isolates. Wilcoxon rank tests: *** for p<0.001; ** p<0.01; * p<0.05.

We focused on JU775 at 25°C, because it allowed for shorter assays, and tested other *E. coli* strains. Culture on the *E. coli* B-type strain BL21(DE3) resulted in a strong Mrt phenotype, while the *E. coli* K-12 derivatives HT115 and C600 partially rescued the phenotype (Figure 5D,E). Mixing a K-12 and a B strain resulted in an intermediate Mrt phenotype (Figure 6A). These results indicated that the Mrt phenotype of wild isolates was revealed when they were cultured on an *E. coli* B-type strain.

**Figure 6.**
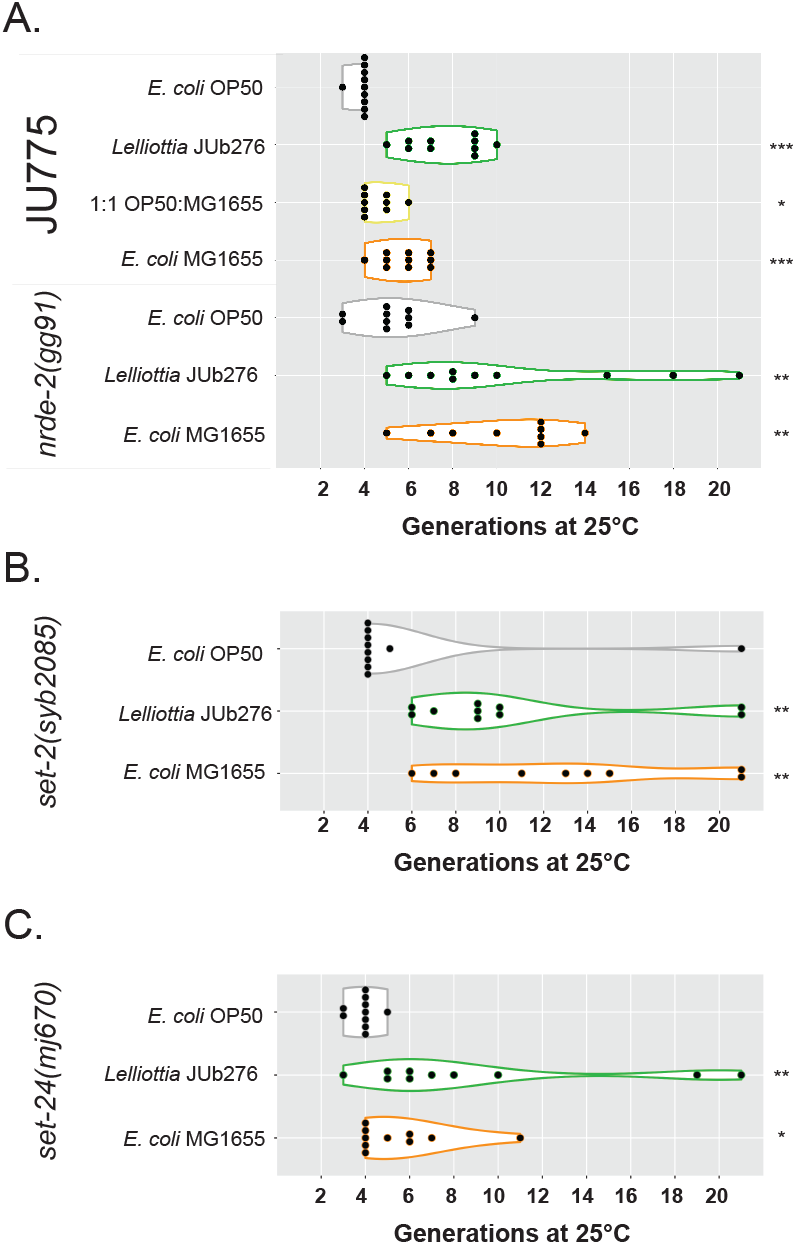
Suppression of the Mrt phenotype of *nrde-2*, *set-2* and *set-24* mutants by a naturally associated bacterium and *E. coli* K-12. (A) *nrde-2(gg91)*, in parallel with JU775. The phenotype of JU775 appears intermediate on a 1:1 mixture of the two *E. coli* strains compared to their pure culture. (B) *set-2(syb2085*). (C) *set-24(mj670)*. Wilcoxon rank tests compared to *E. coli* OP50: *** for p<0.001; ** p<0.01; * p<0.05.

### Mutants in nuclear small RNA pathway or SET-domain genes are rescued by naturally associated bacteria and *E. coli* K-12

We further asked whether the rescue was specific to the Mrt phenotype of wild isolates or also concerned mutants in the N2 reference background with a temperature-sensitive Mrt phenotype.

We tested mutants in two different molecular processes: the *nrde-2* mutant affects the nuclear small RNA response involved in multigenerational silencing inheritance (Burton et al. 2011), while the *set-2* mutation affects a histone H3K4 methyltransferase (Robert et al. 2014; Caron et al. 2022). Both were rescued by the natural associate JUb276 and *E. coli* MG1655 (Figure 6A,B). For *set-2*, the results are consistent with (Robert et al. 2020), who found a suppression by *E. coli* HT115.

We previously showed that the strong Mrt phenotype of the MY10 wild isolate was partially explained by a deletion in the *set-24* gene (Frézal et al. 2018). We tested a *set-24* mutant in the N2 background (courtesy of Giulia Furlan and Eric Miska) and found that its Mrt phenotype was also rescued by the natural bacteria JUb276 and *E. coli* K-12 (MG1655) (Figure 6C).

Thus, we conclude that the Mrt phenotype of laboratory mutants in small RNA inheritance or SET-domain protein genes is suppressed by culture on bacteria other than *E. coli* OP50.

### Multigenerational memory of the bacterial environment

In order to test whether the animals had a memory of their past bacterial environment, we shifted animals of the *C. elegans* JU775 strain cultured for several generations at 25°C on an *E. coli* K-12 strain (on which their Mrt phenotype was suppressed) to *E. coli* OP50 (Figure 7A). We performed this experiment twice, changing the *E. coli* strain (MG1655 or HT115) and the transfer conditions (with or without bleaching at the time of transfer). In both experiments, the lines that were propagated for 10-20 generations at 25°C on an *E. coli* K-12 strain showed a rescued phenotype when transferred back on OP50 (Figure 7). This indicates a multigenerational memory of the bacterial environment.

**Figure 7.**
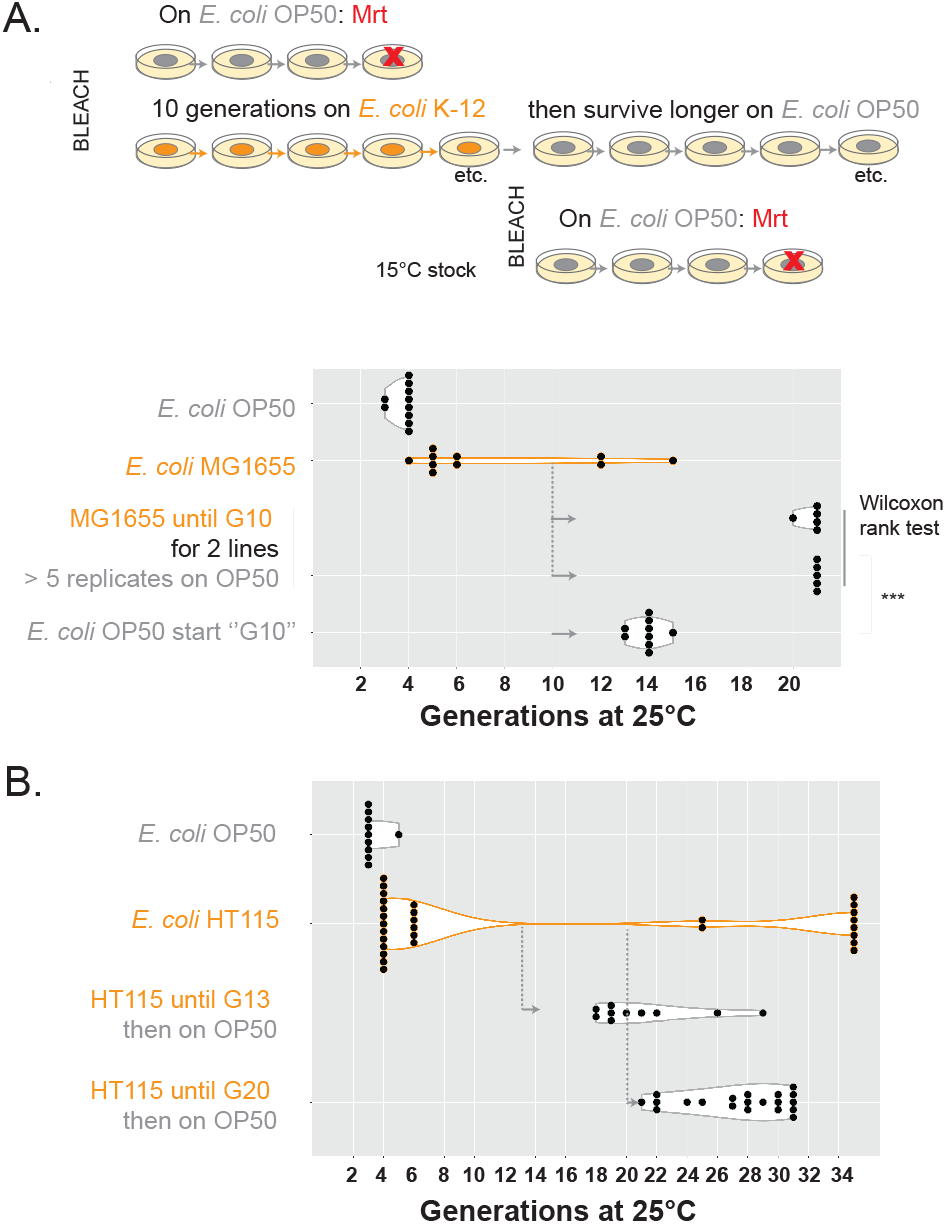
Multigenerational memory of the bacterial environment. In both experiments, the Mrt phenotype of JU775 was assayed on *E. coli* OP50 or a *E. coli* K-12 strain (MG1655 or HT115) at 25°C. After 10, 13 or 20 generations on the K-12 strain, replicates of the surviving lines were maintained on the K-12 strain or transferred back to OP50 (grey arrows). (A) *C. elegans* JU775 with *E. coli* MG1655. Transfer at G10 was with bleaching. For two of the surviving lines, five replicates were assayed. A control from a 15°C culture was bleached in parallel (bottom treatment). Wilcoxon rank test comparing the transferred versus control replicates: *** for p<0.001. (B) *C. elegans* JU775 with *E. coli* HT115 carrying (irrelevant) RNA interference plasmids, as indicated in the Table S2. Transfer was without bleaching, after G13 and G20. A single replicate and two replicates were assayed from G13 and G20, respectively.

## Discussion

### Continuous distribution of the mortal germline phenotype

Expressed as the number of generations to sterility, the Mrt phenotype is a quantitative trait with a continuous distribution (Figure 1). The laboratory reference strain N2 is not fully exempt from the phenomenon and, in our experiments, displayed the sterility phenotype in some replicates (Table S2).

The variance of the Mrt value depends on its mean value (Figure 1). The observed variance distribution makes conditions with intermediate Mrt values particularly difficult to work with. Mrt phenotypes of intermediate strength are intrinsically variable because of the stochastic nature of the trait and its environmental sensitivity. In addition, th phenotype is sensitive to environmental conditions of past generations at 25°C (Figure 7), which further complicates the assays. Despite this experimental caveat, at least some variation in Mrt phenotype among isolates is of genetic origin and can be studied by linkage mapping using laboratory crosses (Frézal et al. 2018) or association in natural populations (here).

### A major GWAS locus underlies this conditionally deleterious variation

The association study detected one major locus on chromosome III. We narrowed down the possible interval to 4.66-6.49 Mb, the association being stronger with the left part (4.66-5.4 Mb). The interval is in a region of low recombination in the center of *C. elegans* chromosomes (Rockman and Kruglyak 2009; Noble et al. 2017), explaining a high linkage disequilibrium and the need for further laboratory recombination. This interval contains a number of candidate polymorphisms, including in germline-expressed genes in which null mutations are known to cause a Mrt phenotype at 25°C: for example, statistically associated polymorphisms include non-synonymous substitutions in *morc-1*, *nhl-2* and *set-2* (Weiser et al. 2017; Herbette et al. 2017; Robert et al. 2020; Davis et al. 2018) and non-coding changes in a number of other candidates (Table S3). The interval also contains many other associated SNPs and insertion-deletion variants (Cook et al. 2017). We caution that methods associated with CRISPR/Cas9 genome editing seemed to produce undesired effects on the Mrt phenotype, at least some of which disappeared after prolonged culture and/or freeze-thawing. Further work will be needed to identify the causal polymorphism(s).

### Germline immortality in natural environmental conditions

In addition to temperature, we detected a strong biotic environmental component to the Mrt phenotype. In particular, the bacterial strain used to feed the nematode had a strong effect. The use of *E. coli* OP50 for *C. elegans* culture dates from the "domestication" of *C. elegans* by Sydney Brenner (Brenner 1974). Many *C. elegans* phenotypes have been recently described to depend on the bacterial culture environment, including the type of *E. coli* strain (Yen and Curran 2016; Zhang et al. 2017; Neve et al. 2020). *E. coli* K-12 and B strains differ in a variety of features, several of which are known to affect other *C. elegans* phenotypes (Jeong et al. 2009; Studier et al. 2009; Reinke et al. 2010; Stuhr and Curran 2020; Daegelen et al. 2009; Pinske et al. 2011; Yoon et al. 2012; Neve et al. 2020; Zhao et al. 2022; Yen and Curran 2016). The use of *E. coli* screens (e.g. Virk et al. 2016; Han et al. 2017a; Qi and Han 2018; Govindan et al. 2019; Zhang et al. 2019; Warnhoff and Ruvkun 2019; Shin et al. 2020) may enable future studies to understand which environmental parameters explain variation in the Mrt phenotype. Importantly, this bacterial influence on the Mrt phenotype suggests that the relevant natural genetic variation may concern its metabolism, immunity or sensory system, in addition or in combination with gene expression regulation by small RNAs and histone modifications. Variation in metabolism and interaction with microbes possibly link to small RNA and histone modification pools (Tauffenberger and Parker 2014; Han et al. 2017b; Nono et al. 2020; Kaletsky et al. 2020; Wan et al. 2022; Legue et al. 2022).

A particular case is that of obligate intracellular intestinal parasites. *N. parisii* and *N. ausubeli* microsporidia are strong suppressors of the Mrt phenotype and hence have a conditional positive effect on population growth. These horizontally transmitted microsporidia are frequently encountered with *C. elegans* (Zhang et al. 2016). They infect the intestine and not the germline, thus showing an effect of the soma on germline maintenance. This constitutes one more example of an absence of a soma-to-germline "Weismann" barrier in *C. elegans* (e.g. Devanapally et al. 2015; Jobson et al. 2015; Rechavi et al. 2014; Moore et al. 2019; Gabaldon et al. 2020).

### The surprising maintenance of a conditional deleterious phenotype

Being strongly deleterious, the Mrt phenotype is likely shielded in the wild, explaining how associated polymorphisms can be maintained at intermediate frequency. The ts-Mrt phenotype can therefore be considered as a consequence of the standard *C. elegans* laboratory culture conditions - useful as a tool to reveal laboratory mutations and natural polymorphisms in small RNA inheritance and histone modification pathways (e.g. *set-24*), as well as to study germline maintenance requirements. Specific environmental conditions may also reveal it in the wild.

Mutation accumulation lines can be used to measure the ease with which a given phenotype can be altered by de novo mutation and, by comparison with natural populations, allow inference of natural selection. With this aim in mind, Saber et al. (2022) used spontaneous mutation accumulation lines in *C. elegans* starting from two wild-derived strains with a non-Mrt phenotype. They found a single mutation line with a strong Mrt phenotype, likely caused by a frameshifting insertion in *nrde-2*, and none with a weak Mrt phenotype. Since many wild isolates have at least a weak Mrt phenotype, they concluded that the Mrt phenotype may be under balancing selection in the wild. Balancing selection is consistent with a varying environmental context of selection of the underlying genetic variation.

The ability to transmit non-genetic information allows the species to transiently provide environmental cues to a variable number of generations. Variation in small RNA inheritance and chromatin (as with *set-24*) may be selected differentially by specific temporal sequences of environmental variation. Beyond the large diversity of Argonaute genes and RNAi mechanisms (Yigit et al. 2006; Winston et al. 2000; Tijsterman et al. 2002; Félix et al. 2011; Paaby et al. 2015; Chou et al. 2022), these nematodes may have evolved variation modulating the memory of non-genetic information.

### Multigenerational sterility

We previously examined cellular features of germline degeneration along the mortal germline process of *C. elegans* wild isolates: germline reduction, defects in chromosome pairing and excess of unresolved double-stranded breaks (Frézal et al. 2018). These cellular features resembles the Mrt phenotype of mutants in small RNA inheritance and histone modifying enzymes, which involves abnormal regulation of gene expression, including repetitive elements (McMurchy et al. 2017; Wallis et al. 2019), as well as genome stability (Herbette et al. 2017) and germline pluripotency (Robert et al. 2014). Their Mrt phenotype appears however distinct from that of the *prg-1/piwi* mutant (for which the cause of sterility is debated; Barberan-Soler et al. 2014; McMurchy et al. 2017; Heestand et al. 2018; Barucci et al. 2020; Reed et al. 2020; Wahba et al. 2021; Spichal et al. 2021): especially, the latter does not show temperature dependence and is suppressed by starvation (Simon et al. 2014)..

The temperature-dependent Mrt phenotype itself is a manifestation of memory of the temperature environment across generations. Whether this temperature dependence reflects a variation in metabolism, germ granules, chromatin compaction, nucleic acid hybridization or any other process remains to be investigated.

In addition to the cumulative nature of the Mrt phenotype over generations, we demonstrated that a wild *C. elegans* isolate keeps a further multigenerational memory of the bacteria on which the strain was cultured at 25°C. This suggests that its Mrt phenotype is not only due to a metabolic requirement during the tested generations but may result from metabolic carryover of the physiological or epigenetic state of the animals. Such memory has been observed in *C. elegans* using various experimental paradigms (Baugh and Day 2020; Rechavi et al. 2014; Jobson et al. 2015; Gabaldon et al. 2020; Moore et al. 2019; Ma et al. 2019; Zhang et al. 2021). Further work will be necessary to study its molecular basis. A possible mechanism involves changes in the pools of small RNAs and histone modifications (Dodson and Kennedy 2019), or their metabolic precursors. Alternatively, other processes, such as the quality or quantity of mitochondria (Zhang et al. 2021), may be at play in this multigenerational environmental memory.

## Material and Methods

### *C. elegans* culture and strains

*C. elegans* was cultured on 55 mm NGM agar plates (Stiernagle 2006) seeded with 100 μl of a saturated culture of *E. coli* OP50 grown overnight in LB at 37°C, except if otherwise indicated. *C. elegans* wild isolates for the association analysis were obtained from CGC, Michael Ailion, Erik Andersen, Cori Bargmann, Christian Braendle, Elie Dolgin and Asher Cutter, Matt Rockman, Paul Sternberg or our own collection. The *set-2(syb2085)* mutant (Caron et al. 2022) was a gift of Francesca Palladino and the *set-24(mj670)* mutant of Giulia Furlan and Eric Miska. The *nrde-2(gg91)* mutant (Burton et al. 2011) was obtained from CGC.

Bleaching was performed by spotting gravid adults or embryos onto an NGM plate seeded with *E. coli* OP50 and immediately adding 50 μl of the bleach/NaOH solution (Stiernagle 2006).

### E. coli strains

OP50 was acquired from Paul Sternberg’s lab in 1994. MG1655 was received from Javier Apfeld’s lab, C600 from John Chapman’s lab, BL21(DE3) from Bertrand Ducos (ENS). HT115 was acquired from Andy Fire and transformed with the L4440 vector carrying various inserts, as indicated in Table S2 (Timmons et al. 2001).

### Mortal germline assays

Mortal germline (Mrt) assays were carried out as in (Frézal et al. 2018), transferring three L4 stage animals at each generation onto a new plate. Before the start of the assay, the strains were maintained at 15°C with food for two or more generations prior to shifting at the higher temperature. The assays were performed at 25°C, unless otherwise indicated. With this method, animals are never starved. We recorded the generation *n* of culture at 25°C at which no larva was produced (Generation *n* or *Gn*). The assays were generally stopped once the fertility of the 20^th^ generation was scored (reported as G21 in the graphs).

### Genome-wide association analysis

We used data from several Mrt assays performed in different laboratories across several years (Table S2). Each Mrt assay was conducted in a number of replicates, from 1 up to 6. Each assay represents a block, indicated by the date, experimenter and the laboratory where the assay was performed. In total we studied 12 blocks, performed by four experimenters: Marie-Anne Félix (MAF), Lise Frézal (LF), Emilie Demoinet (ED, data from Frézal et al. 2018) and Sayran Saber (SS, data from Saber et al. 2022). The experiments were conducted in four different laboratories: ours in Paris, Eric Miska’s lab in England, Christian Braendle’s in Nice and Charles Baer’s lab in Florida.

To plot the distribution of the Mrt-value across the dataset (Figure 1B), the mean Mrt-value for each strain was calculated as the mean of all the Mrt-values of any experiment where the strain was assayed.

For the GWAS, the strains that were fertile after 20 generations were considered non-Mrt and we set their Mrt-value at 30 generations. We removed the outlier Bergerac strain derivative RW7000 that was sterile at the first generation due to a temperature-sensitive *zyg-12* mutation that is unrelated to the Mutator phenotype of some of its derivatives (Fatt and Dougherty 1963; Mori et al. 1988; Malone et al. 2003; Nigon and Félix 2017; Frézal et al. 2018).

We used the lme4 package (Bates et al. 2015) in R to fit generalized linear mixed-effect models using a Poisson distribution in Rstudio version 1.2.5001 (Team 2015). Block, which is confounded with experimenter, was significantly associated with phenotype (deviance = 3304.0 against 3451.7 for the reduced model with isotype as the sole random effect). For GWAS we extracted isotype BLUPs (best linear unbiased predictions) using the ranef function on the full model: glmer(value∼ (1|block) + (1|isotype), Mrt_ALL, family="poisson"). Residuals were approximately normal (Shapiro-Wilk W=0.988).

To perform the GWAS, we used the SNPs from the 2021-01-21 version of the hard-filtered vcf at CeNDR (Cook et al. 2017), which we filtered using the PLINK toolset (https://zzz.bwh.harvard.edu/plink/). We took all SNPs with a minimum allele frequency of 0.05, and removed markers in complete linkage disequilibrium with the indep-pairwise command using a sliding window size of 100,000 bp, a count of 50,000 variant per window and a maximum r^2^ of 0.99. We performed further GWAS on specific regions on chromosome III (from 4 to 10 Mb), using all SNPs from the soft-filtered vcf from CeNDR.

We used the GridLMM package to test SNP associations with the Mrt BLUPs (Runcie 2018), controlling for population structure with an additive kinship matrix. The significance threshold was calculated from 1000 phenotype permutations (using the *permuts* function, code in Annex).

### Introgressions

All crosses were performed at 18°C. To produce the introgression lines, the chromosome III of JU775 was introgressed into the N2 background by a minimum of six rounds of backcrossing followed by marker genotyping on the different chromosomes (see primers and genotypes in Table S1). Two independent introgressions were made, creating the strains JU3194 and JU3195.

The *mjIs31*[*pie-1::gfp::h2b*] transgene was initially introduced by others using an artificial Mos1 transposon insertion site at 8.4 Mb on chromosome II of the N2 reference strain (Ashe et al. 2012), creating the strain SX461. For another project, we introduced this *mjIs31*[*pie-1::gfp::h2b*] transgene into the above introgressed strain JU3195 via two independent crosses, creating the strains JU3394 and JU3395.

To use as a control, this transgene was backcrossed six times into the wild isolate JU775, creating the independent strains JU3306 and JU3307 (Table S1). (Although *set-24* may be irrelevant in the case of JU775, we took care that these strains carry JU775 alleles in the region of this gene, which carries a partial deletion in MY10 and is located on the same chromosome as the *mjIs31*[*pie-1::gfp::h2b*] transgene).

### Generation of recombinants

Recombination within chromosome III was performed by crossing the males of the introgressed lines JU3395 (JU775 chrIII>SX461) to SX461 hermaphrodites. The crosses were performed at 20°C and the following generations were kept at 18°C to prevent selection of the non-Mrt alleles. For each cross, one F1 heterozygote hermaphrodite was selected randomly and about 100 F2 individuals were singled and genotyped for a recombination event between two markers on chromosome III at 5.4 and 11.0 Mb (Table S1). We then selected recombinant lines that were homozygous at both sites. To this end, we singled 8 F3 animals from the lines homozygous at one site but heterozygous at the other site; we genotyped these F3 animals and selected one with a homozygous recombinant genotype. Multiple recombinant lines were thus obtained: JU4312, JU4336-41, JU4348.

### Microbe isolation and rDNA sequencing

New *C. elegans* isolates and their associated microbes were collected from natural substrates as described in (Félix and Duveau 2012; Barrière and Félix 2014). JU3224-3228 were isolated from samples of rotting vegetal matter from New Zealand, 1-3 weeks after sample collection in April 2017. JU4134 was isolated from the decaying stem of a plant of the Asteraceae family in Santeuil (France) on October 30, 2020. In this case, the sample was placed on a NGM plate with *E. coli* OP50 a few hours after collection. *C. elegans* is generally fully homozygous at all loci in natural populations (Barrière and Félix 2005; Richaud et al. 2018); to make sure our lines were isogenic, strains were selfed by isolation of a single individual for 4-5 generations before further use.

To assess the microbial contribution to the Mrt phenotype of JU4134, a procedure via a microbial preparation was used because the experimenter did not expect that single bacteria could rescue the Mrt phenotype and wished to go stepwise from the most ecologically relevant environment. A microbial preparation was first made by incubating the nematodes overnight with 10 mM tetramisole in 15 ml tubes, with mild mixing on a rotator. The nematode corpses were pelleted by centrifugation at 1000 g. The preparation was used for seeding NGM plates or for further isolation of bacteria. This tetramisole procedure did not kill all nematodes so seeded plates were inspected after a few days and discarded if they contained a nematode that had survived.

Microbial strains JUb277-283 were then isolated from the JU4134 microbial preparation by streaking on 90 mm NGM plates without bacteria. Individual colonies were further isolated by streaking on a new plate. JUb283 was isolated directly from the JU4134 culture. Bacteria showing different colony and cell aspects when observed with the dissecting microscope or under Nomarski optics were retained. The bacteria were given a JUb identifier.

The bacterial strain JUb276 was isolated from the unbleached culture of JU3224 by streaking the microbial culture surrounding the nematodes onto 90 mm NGM plates. The plates were incubated at room temperature. The streaking procedure was repeated to isolate single colonies. One colony was then used as JUb276 and a culture derivative was frozen.

PCR primers for bacterial rDNA sequencing are in Table S1. NCBI accession numbers are ON110486-90. Sequences of JUb277 and JUb282 were identical, as were those of JUb279, JUb280 and JUb283.

Bacteria were grown overnight in liquid LB with shaking, at 37°C for *E. coli*, at 25°C for the other bacteria. A 100 μl drop of the culture was then spotted in the center of a 55 mm NGM nematode culture plate (50 μl in some experiments, as indicated in Table S2). The plates were left at room temperature for 2-3 days and stored in a cold room before use. Care was taken to equilibrate the plates at the relevant temperature before use for the Mortal germline assay.

### Mortal germline assays in the presence of microbes

Several methods were used to expose nematodes to microbes during the Mortal germline assays; to each corresponds a colour code in the figures. The assay could be performed with the original microbial fauna (maroon), before the *C. elegans* culture was bleached. All other methods started with a bleached nematode culture. In the first case, the plates were seeded with the microbial tetramisole preparation (see above), in the absence of *E. coli*, and the nematode culture was bleached on this plate. In the second method (green), the plate was seeded by one or several microbial strains isolated as above, cultured overnight in LB at 25°C in a shaker. The latter method was also used for culture on different *E. coli* strains.

The *C. elegans* stock was maintained at 15°C on OP50 without starvation, and for two-three generations with the relevant bacteria at 15°C, before the mortal germline assay was started by shifting to 25°C (see details corresponding to each experiment in Table S2).

Microsporidial spores were prepared and quantified as in (Zhang et al. 2016). 20 uninfected young adults (i.e. prior to first egg formation) were transferred to a 55 mm NGM plate seeded with *E. coli* OP50. 5 million microsporidian spores in 100 μl distilled water were placed onto the *E. coli* lawn. The cultures were then incubated at 15 °C. The infection symptoms were checked by Nomarski optics at 72 hours after inoculation. Ten L4 larvae from the infected population were transferred to a fresh plate and moved to 23°C (Generation 1) and then ten L4 animals were transferred onto fresh plates at each generation (every 2-3 days, generally 3 transfers per week). Uninfected individual L4 larvae were also tested in parallel. We used 23°C because the detrimental effects of the microsporidia were too high at 25°C. In this experiment, the transfer was stopped when 10 L4 stage animals were not available.

*E. coli* HT115 bacteria carrying (irrelevant) RNA interference plasmids, and OP50, were grown overnight in liquid LB with shaking, at 37°C, and then seeded (100 μl) onto NGM plates supplied with 1 μM IPTG. Before the start of the assay, the *C. elegans* strains were maintained at 15°C on OP50. To initiate the Mrt assays, five gravid hermaphrodites were spotted onto freshly prepared IPTG-plates (24-32 hrs), immediately adding 50 μl of the bleach/NaOH solution (Stiernagle 2006) before incubating plates at 25°C. The following steps are identical as the Mortal germline assays described above, transferring three L4 stage animals at each generation onto a new plate. In order to assess the memory of past bacterial environment, at G13 and G20, for all the fertile lineages, three L4 stage animals were transferred back to OP50 seeded plates at 25°C, the following steps are similar to the Mrt assay with the transfer of three L4 stage animals at each generation onto a new OP50 seeded plate.

### Statistical analysis of Mrt assays

The Mrt trait quantified as the number of generations to sterility has a low variance when the number is small, a high variance when it is larger and again a low variance when none of the replicates become sterile by 20 generations. We thus used a conservative rank test for statistical testing, specifically the wilcox.test in R. We only compared assays run in parallel, represented in a given figure panel.

## Acknowledgements

We are grateful to Hervé Gendrot for pouring large numbers of agar plates for this project. We thank Nathalie Pujol and Jonathan Ewbank for sending samples from New Zealand; Giulia Furlan and Eric Miska for the unpublished *set-24* CRISPR mutant and Francesca Palladino for the *set-2* CRISPR mutant. We thank Javier Apfeld’s lab for sending MG1655 and John Chapman’s for C600. Some strains were provided by the CGC, which is funded by NIH Office of Research Infrastructure Programs (P40 OD010440). We thank WormBase and CeNDR. This work was funded through grants from the Agence Nationale de la Recherche ANR-14-CE10-0003 and ANR-19-CE12-0025. M.S. was supported by a PhD fellowship from the French Ministry of Research and the Fondation pour la Recherche Médicale PhD FRM FDT202106013206.

## Supplementary Material

**Figure S1. GWAS using a broader panel including divergent strains from the Pacific region.** The top panel includes the isolates in the red and yellow groups in Crombie et al. (2019) but not the more divergent Pacific strains. The middle panel includes all 132 isolates. The bottom panel shows the absence of correlation of the Mrt-value with temperature at the time of collection, using all 132 isolates.

**Figure S2. Additional experiments during isolation of *C. elegans* wild isolates and their associated microbes**. (A) The five C. *elegans* strains that were isolated from New Zealand are shown. JU3224 showed the clearest rescue of the Mrt phenotype by associated microbes and was used for isolation of a bacterial clone. (B) Intermediate steps in the isolation of bacteria from JU4134. The top panel shows the effect of adding a levamisole-treated microbial preparation of the JU4134 culture, a mix of isolated bacteria and eucaryote, or pairs of bacteria. The experiment in the top panel is shown partially in Figure 3C.

**Figure S3. Starvation does no strongly rescue the Mrt phenotype of the wild isolate JU775.** The two graphs represent two experiments performed successively. The “continuously fed” controls were started from a 15°C culture at different times. The first experiment was performed in parallel to that in Figure 5C. The second experiment serves as a control for the 19-day starvation treatment of the first experiment, fed at the same time as the second experiment was started at 25°C. The pairing of fed and starved lines is indicated in Table S2.

**Table S1. Strains and genotyping primers.**

**Table S2. Mortal germline assays.**

**Table S3. Finer GWAS for the set of 115 strains with all soft-filtered SNPs from CeNDR between 4 to 7 Mb on chromosome III.**

