## Supplementary material for "Genome-wide association and environmental suppression of the mortal germline phenotype of wild *C. elegans*": Figure S1

Genome Wide Association Study on a subset of 126 wild isolates

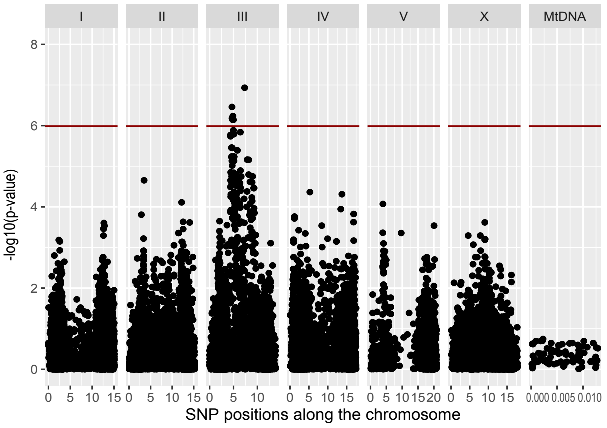


Genome Wide Association Study on a subset of 132 wild isolates

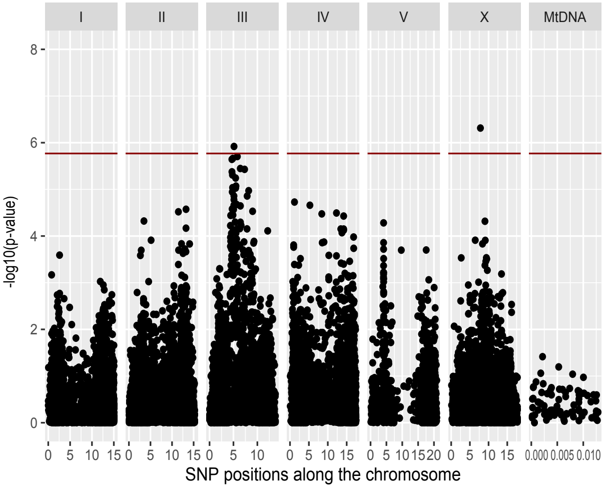


Mrt-value of 132 wild isolates in function of the temperature at sampling location

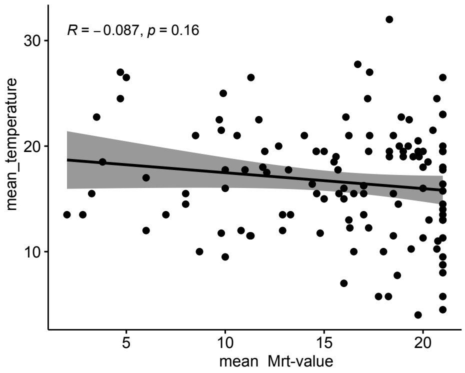


Figure S1
