## Supplementary figures and images for "Genome-wide association and environmental suppression of the mortal germline phenotype of wild *C. elegans*"

### Figure S2

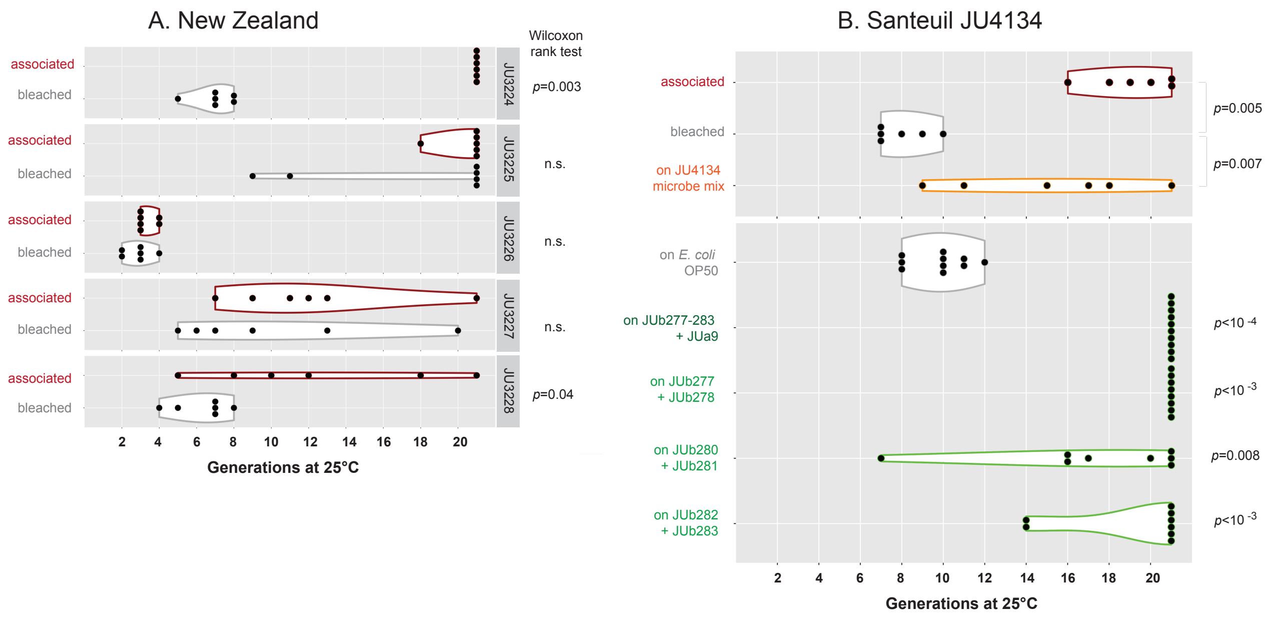

### Figure S3

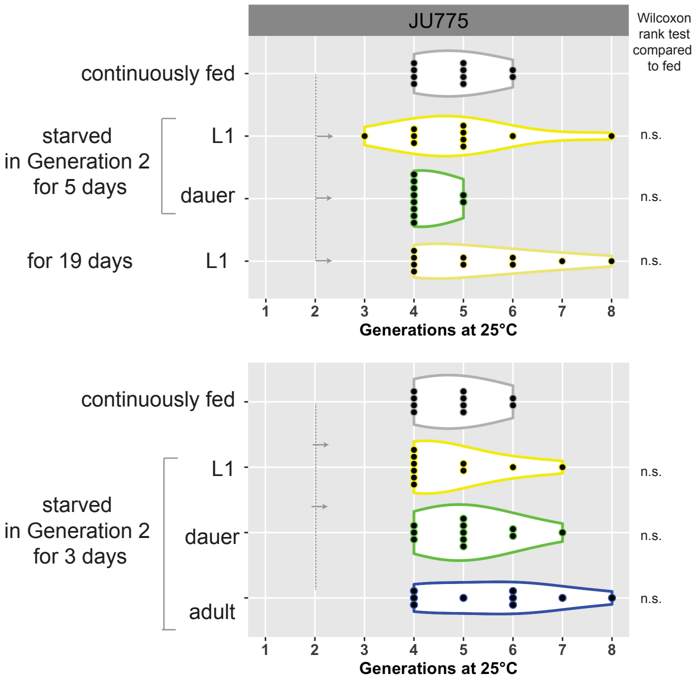
